# Iron Inhibits Glioblastoma Cell Migration and Polarization

**DOI:** 10.1101/2022.10.13.512175

**Authors:** Ganesh Shenoy, Sina Kheirabadi, Zaman Ataie, Kondaiah Palsa, Quinn Wade, Chachrit Khunsriraksakul, Vladimir Khristov, Becky Slagle-Webb, Justin D. Lathia, Hong-Gang Wang, Amir Sheikhi, James R. Connor

**Author notes:** **Corresponding Author:** James R. Connor, Ph.D. Distinguished Professor and Vice-Chair of Research, Department of Neurosurgery, Penn State College of Medicine, Hershey, PA, 17033.

## Abstract

Glioblastoma is one of the deadliest malignancies facing modern oncology today. The ability of glioblastoma cells to diffusely spread into neighboring healthy brain makes complete surgical resection nearly impossible and contributes to the recurrent disease faced by most patients. Although research into the impact of iron on glioblastoma has addressed proliferation, there has been little investigation into how cellular iron impacts the ability of glioblastoma cells to migrate - a key question especially in the context of the diffuse spread observed in these tumors. Herein, we show that increasing cellular iron content results in decreased migratory capacity of human glioblastoma cells. The decrease in migratory capacity was accompanied by a decrease in cellular polarization in the direction of movement. Expression of CDC42, a Rho GTPase that is essential for both cellular migration and establishment of polarity in the direction of cell movement, was reduced upon iron treatment. Bioinformatic analysis of CDC42 mRNA revealed a potential iron-responsive element that may contribute to the regulation of CDC42 by iron. We then analyzed a single-cell RNA-seq dataset of human glioblastoma samples and found that cells at the tumor periphery had a gene signature that is consistent with having lower levels of cellular iron. Altogether, our results suggest that cellular iron content is impacting glioblastoma cell migratory capacity and that cells with higher iron levels exhibit reduced motility.

## 1. Introduction

Glioblastoma remains one of the most difficult to manage malignancies facing modern medicine with five-year survival rates being only around 6% [1]. One of the reasons glioblastoma confers such a dismal prognosis is due to the ability of glioblastoma cells to migrate and spread into neighboring healthy brain tissues, making complete surgical resection nearly impossible [2,3]. Consequently, nearly all patients suffer from recurrent disease despite undergoing a rigorous treatment regimen with maximal surgical resection, chemotherapy, and radiation therapy [4,5]. Given the essential role glioblastoma cell migration plays in contributing to recurrent disease, it is crucial to understand the processes underlying dissemination of glioblastoma cells into healthy brain.

Iron is a nutrient that is known to be essential for cellular growth and function. Numerous enzymes, including those involved in DNA and RNA synthesis, utilize iron as a cofactor [6]. The role of iron in cancer cells is particularly a topic of interest given the necessity of iron for cellular proliferation [7–10]. Additionally, iron generates reactive oxygen species (ROS) and activates downstream signaling pathways associated with redox homeostasis, some of which promote tumor progression [7–10]. While much work has focused on the role iron plays in cellular proliferation and ROS generation, little is known regarding iron’s role in cell migration in glioblastoma.

Here we characterize how iron impacts cell migration in T98G and LN229 human glioblastoma cells. We demonstrate that iron inhibits cell migration in both cell lines and that this inhibition is accompanied by a decrease in the expression of CDC42, a Rho GTPase that is essential for appropriate cellular polarization [11,12]. We find that the iron-induced decrease in CDC42 is accompanied by functional reduction of polarization in migrating cells. We also show through analysis of a single-cell RNAseq dataset that cells in the tumor periphery (i.e., the cells spreading into healthy brain) appear to express a gene pattern associated with decreased iron levels – consistent with our finding that iron inhibits cell migration in glioblastoma cells.

Our findings describe an understudied aspect of iron in glioblastoma cell biology – its effect on cell migration. Moreover, our findings provide evidence for further evaluation of how iron impacts patient outcomes in glioblastoma. Given the increased interest in the use of small paramagnetic iron oxide nanoparticles for tumor imaging, understanding the impact of iron on the behavior of glioblastomas is critical [13,14].

## 2. Materials and Methods

### 2.1 Cell Culture

T98G (ATCC, Catalog #: CRL-1690) and LN229 (ATCC, Catalog #: CRL-2611) authenticated human glioblastoma cell lines were purchased from the American Type Culture Collection (ATCC). T98G cells were cultured in RPMI 1640 supplemented with GlutaMAX™ (ThermoFisher Scientific, Catalog #: 61870127), 10% fetal bovine serum (GeminiBio, Catalog#: 100-106), and 1% Penicillin-Streptomycin (ThermoFisher Scientific, Catalog #: 15140-122). LN229 cells were cultured in DMEM supplemented with GlutaMAX™ (ThermoFisher Scientific, Catalog #: 10567022) with 10% fetal bovine serum, and 1% Penicillin-Streptomycin. Cells were maintained in a humidified 5% CO2 tissue culture incubator at 37 °C and were used within 10 passages post-thawing to maintain reproducibility.

### 2.2 Wound Healing Assays

An Incucyte® Live-Cell Imaging System was used to study migration in T98G and LN229 glioblastoma cell lines. 5 x 10^4^ cells were seeded in each well of an Incucyte® ImageLock 96-well plate (Sartorius, Catalog #: 4379) in 100 μL/well of either complete RPMI or DMEM, respectively, and allowed to adhere to the plate for 24 hours. Cells were then scratched using an Incucyte® Woundmaker Tool (Catalog #: 4563) and washed with phosphate-buffered saline to remove cell debris. 0 – 300 μM ferric ammonium citrate (FAC) (Sigma-Aldrich, Catalog #: F5879) was added and phase contrast images were acquired in 2-hour intervals. The relative wound density (RWD) was then quantified as a function of time to determine migratory capacity. The relative wound density is a measure of the cell density in the wounded area relative to the cell density in the area outside the wounded area and provides normalization for alterations in cell density caused by proliferation [15]. For deferoxamine rescue experiments, the migration studies were carried out as described above but with or without the addition or 0 – 200 μM deferoxamine to the cells.

### 2.3 Three-dimensional (3D) Granular Hydrogel Cell Migration Assays

3D granular hydrogel scaffolds were synthesized from gelatin methacryloyl (GelMA) hydrogel microparticles as previously described [16–18]. Further details regarding the hydrogel scaffold synthesis are available in the Supplementary Information. To evaluate cell migration, all cells were labeled with CellTracker™ Green 5-chloromethylfluorescein diacetate (CMFDA) Dye based on the manufacturer protocol, followed by topical cell seeding on 3D microgel scaffolds. Prior to the seeding, scaffolds were soaked in DPBS supplemented with 1% v/v antibiotic for 3 h in a 24-well plate non-treated cell culture plate. Then, DPBS was removed from the plate, and 20 μL of stained cell suspension (5×10^6^ cells per mL of FAC-treated or untreated culture media) was added to each scaffold. Scaffolds were incubated for 30 min, permitting cells to attach to the microgels, followed by adding complete culture media and incubation at 37 °C in a CO2 atmosphere (5% v/v) for 72 h. Finally, samples were cross-sectioned and imaged using a Leica DMi8 THUNDERED™ microscope (Germany), with an excitation wavelength of 470 nm (blue) and an emission wavelength of 550 nm (green). All images were analyzed using the ImageJ software (FIJI, version 1.53c, NIH, MD, USA) [19] and statistics were computed using Mathematica. One-way analysis of variance (ANOVA) test, with subsequent Tukey and Bonferroni post hoc tests, were performed to analyze statistical significance.

### 2.3 Cell Proliferation Assays

Cell proliferation was monitored using both CyQuant™ cell proliferation assays (ThermoFisher Scientific, Catalog #: C7026) as well as optical confluence measurements. For CyQuant™ cell proliferation assays, 3,000 T98G or LN229 cells were seeded in 96 well plates and allowed to adhere overnight. Cells were then treated with 0 – 400 μM FAC (Sigma-Aldrich, Catalog #: F5879) and measurements were collected at t = 0, 24, 48, and 72 hours post treatment following manufacturer instructions. Briefly, treatment media was removed from each well and plates were frozen at −80°C. Plates were then thawed and CyQuant reagent was added to the plates. Fluorescent measurements were then collected using a plate reader with excitation at 480 nm and emission at 520 nm. For optical confluence measurements, 3,000 T98G or LN229 cells were seeded in an Incucyte® ImageLock 96-well plate and allowed to adhere for 24 hours. Cells were then treated with 0 – 400 μM ferric ammonium citrate and optical confluence measurements were acquired in two-hour intervals and automatically quantified using Incucyte® software.

### 2.5 Cell Polarization Assay

Cell polarization assays were performed as described previously [20,21]. Briefly, 5 x 10^4^ T98G or LN229 cells were seeded in each well of a 96-well plate and allowed to adhere to the plate for 24 hours. Cells were then scratched using an Incucyte® Woundmaker Tool (Catalog #: 4563) and washed with PBS with calcium and magnesium to remove cell debris. Cells were then treated with 0 – 200 μM FAC or 0 – 100 μM hemin and cells were allowed to migrate for 6 hours. The migrating cells were then fixed with 4% formaldehyde (ThermoFisher Scientific, Catalog #: 28908) and subsequently permeabilized in 0.2% Triton-X-100 (Sigma-Aldrich, Catalog#: X100). Cells were then stained with anti-pericentrin (Abcam, Catalog #: ab220784) and DAPI (ThermoFisher, Catalog #: 62248) and visualized using a fluorescent microscope. Cells were classified as correctly polarized if the centrosome fell within the quadrant facing the scratched area relative to the cell nucleus. Cells in which more than one centrosome was observed were scored as positive if at least one of the centrosomes fell within the quadrant facing the scratched area relative to the cell nucleus. The individual scoring cells was blinded to experimental conditions to reduce bias.

### 2.4 Immunoblotting

Roughly 3.5 x 10^5^ T98G or LN229 cells were seeded in 6-well plates (Corning Life Sciences, Catalog #: 353046) and allowed to adhere for 24 hours. The cells were subsequently treated with 0 – 200 μM FAC or 0 – 100 μM hemin (Sigma-Aldrich, Catalog #: 51280-1G) for 6 hours. The treatment media was then removed from each well and cells were washed with cold PBS with calcium and magnesium. Cells were then lysed in 2X Laemlli buffer (Bio-Rad, Catalog #: 1610737) supplemented with 5% β-mercaptoethanol (ThermoFisher Scientific, Catalog #: 21985023) and then agitated using a vortexer set at 1000 RPM for 30 minutes. 20 μL of cell lysates were then loaded into precast 4-20% polyacrylamide SDS-PAGE gels (Bio-Rad, Catalog #: 5671094) for electrophoretic separation. Proteins were then transferred onto PVDF membranes (Cytiva, Catalog #: 10600023) using a wet transfer approach and membranes were then blocked with 3% bovine serum albumin (Tocris, Catalog #: 5217) and probed for CDC42 (Cell Signaling Technology, Catalog #: 2466S) and β-actin (Sigma-Aldrich, Catalog #: A5441).

### 2.6 Measurement of Labile Iron Pool with SiRhoNox-1

The cell labile iron pool was assessed using SiRhoNox-1 (Sigma-Aldrich, Catalog #: SCT037), a dye that becomes irreversibly fluorescent in the far-red spectrum upon binding specifically to Fe^2+^. Roughly 5 x 10^4^ T98G or LN229 cells were seeded in 96 well plates and allowed to adhere for 24 hours. The cells were then treated with 0 – 200 μM FAC or 0 – 100 μM hemin for 6 hours. Cells were then stained with SiRhoNox-1 following manufacturer instructions. Briefly, cells were washed with serum-free media and incubated with 5 μM SiRhoNox-1 in serum-free media for 1 hour at 37 °C in a tissue culture incubator. Unbound dye was then removed by washing cells with PBS with calcium and magnesium and cells were then imaged using a fluorescent microscope with the Cy5 filter set. Fluorescent intensity was subsequently quantified for each image using ImageJ software.

### 2.7 Analysis of Single Cell RNASeq Data

Single cell RNASeq data of 1091 neoplastic cells that were collected between tumor core and periphery locations from four glioblastoma patients by Darmanis et al (GSE84465) was downloaded from gbmseq.org [22]. The authors’ annotations of cell type and location were used to differentiate neoplastic cells isolated from the tumor core and tumor periphery. Gene expression of genes correlated with the iron status of cells (TFRC, FTH1, and FTL) were compared between neoplastic cells in the tumor core and tumor periphery. Statistical computing was performed using R Studio (version 1.2.5033) running R version 3.6.3.

### 2.8 Data Availability

The data generated within this study is available upon request to the corresponding author. Single-cell RNASeq data that was analyzed in this study was obtained from gbmseq.org and is also available on the Gene Expression Omnibus (GSE84465).

## 3. Results

### 3.1 Iron Inhibits Cell Migration in T98G and LN229 Glioblastoma Cells in Wound Healing Assays

We utilized an Incucyte® Live-Cell imaging system to study how iron impacted glioblastoma cell migration. Scratching confluent monolayers of T98G or LN229 cells followed by treatment with 100 – 300 μM iron in the form of ferric ammonium citrate (FAC) showed a significant decrease in cell migration in iron-treated T98G and LN229 human glioblastoma cells compared to non-treated control groups as assessed by live-cell imaging for 24 hours post-scratching (**Figure 1A-D**). The analysis was limited to 24 hours to minimize confounding by cell proliferation. Additionally relative wound density (RWD), a ratio of cell density within the wound relative to cell density outside the wound, was used as the measure of migratory capacity. Using relative wound density as a measure of cell migratory capacity minimizes the impact of proliferation since it also accounts for cell density outside the wound [15]. Interestingly, the iron-induced inhibition of migration appeared to be saturatable with 200 and 300 μM iron concentrations producing similar levels of inhibition of migration.

**Figure 1.**
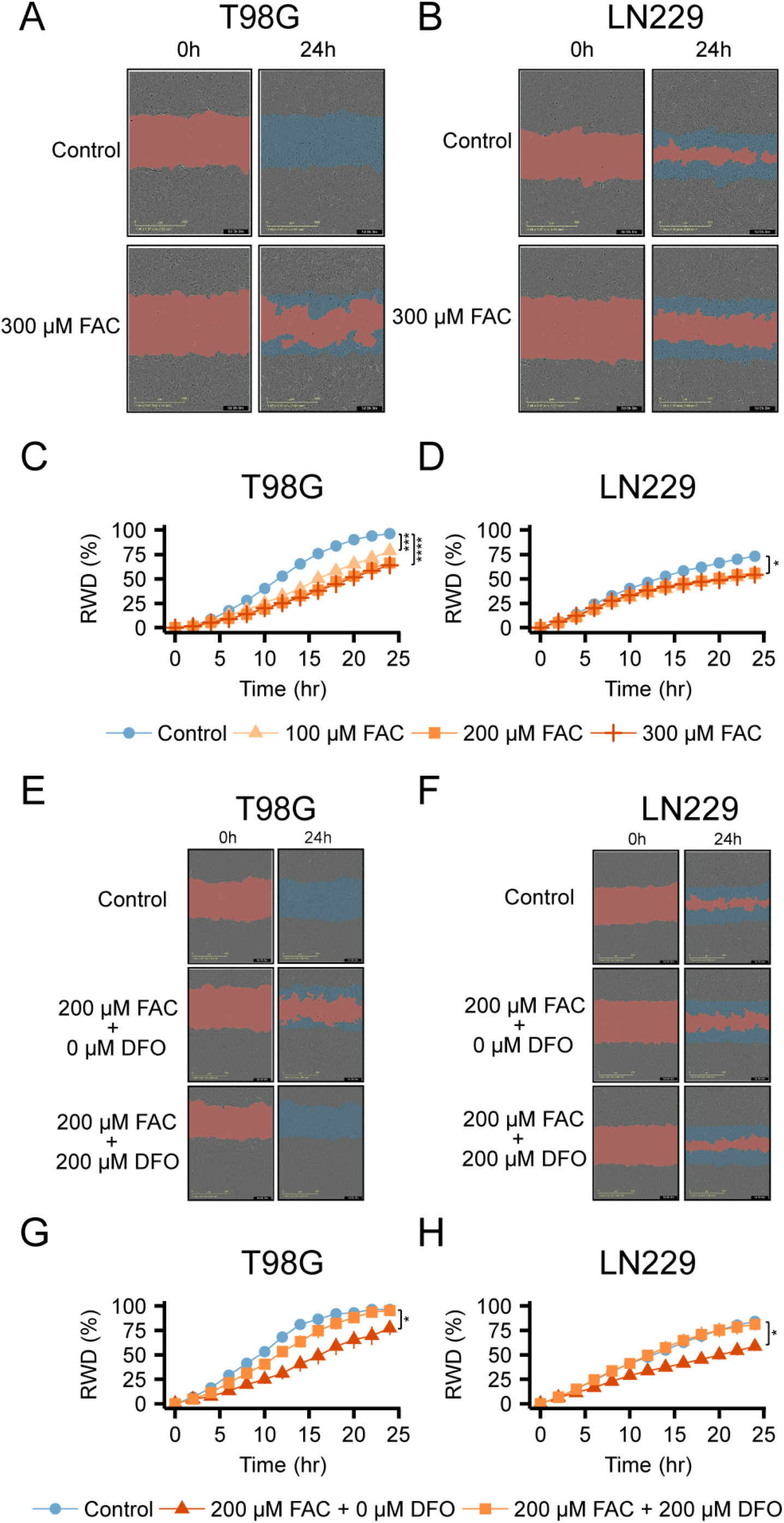
Iron inhibits cell migration in T98G and LN229 cells. Confluent monolayers of **(A)** T98G or **(B)** LN229 glioblastoma cells were scratched to generate cell-free zones and cells were allowed to migrate in the presence of 0 – 300 μM ferric ammonium citrate (FAC). Relative wound density was quantified as a function of time **(C, D)** which revealed that iron inhibited cell migration in these cells. The addition of equimolar concentrations of deferoxamine (DFO), an iron chelator that chelates iron in a 1:1 molar ratio, resulted in rescue of the iron-induced inhibition of cell migration in both **(E, G)** T98G and **(F, H)** LN229 cells. Results representative of at least n = 3 replicates. Student’s two-tailed t-test: *: p < 0.05, **: p < 0.01, ***: p < 0.001, ****: p < 0.0001.

To verify that iron was responsible for the inhibition of migration observed in the wound-healing assays, we performed rescue experiments using deferoxamine, an iron chelator that chelates iron at a 1:1 molar ratio [23]. In wound healing experiments where 200 μM FAC was used to inhibit cell migration, the addition of equimolar concentrations of deferoxamine was able to largely rescue the iron-induced inhibition of cell migration, suggesting that the inhibition of cell migration observed in FAC treated cells is truly due to iron (**Figure 1E-H**).

### 3.2 Iron Inhibits Migration of T98G and LN229 Cells in 3D Granular Hydrogel Scaffolds

To test if the iron-induced reduction in migration observed in the wound-healing assays also occurred during 3D migration, the effect of iron on cell migration was analyzed using a microporous 3D granular hydrogel scaffold, partially mimicking the native brain tissue microenvironment. To this end, T98G and LN229 cells were labeled with a fluorescent dye, followed by 3D topical seeding on granular hydrogel scaffolds (**Figure 2A)**. Granular hydrogel scaffolds provide interconnected microporosity, enabling oxygen, nutrient, and cellular waste transport [16,24]. The microporosity enables nutrient gradient generation, facilitating cell migration within the scaffold [25]. **Figure 2B** presents microscopy images of cross-sectioned scaffolds 72 h after cell seeding. As in the 2D wound-healing assays, iron treatment inhibited migration in both cell lines (**Figure 2C)**. Interestingly, like the 2D migration assays, the decrease in average migration length of FAC-treated cells appeared saturatable and did not significantly decrease at FAC concentrations beyond 100 μM. To evaluate the effect of FAC concentration on cell migration independent of initial cell penetration due to the porous nature of scaffolds, results were compared with average initial cell penetration length (*t* ~ 4 h), where cell movement is governed by gravitational and capillary forces (**Figure 2C**).

**Figure 2.**
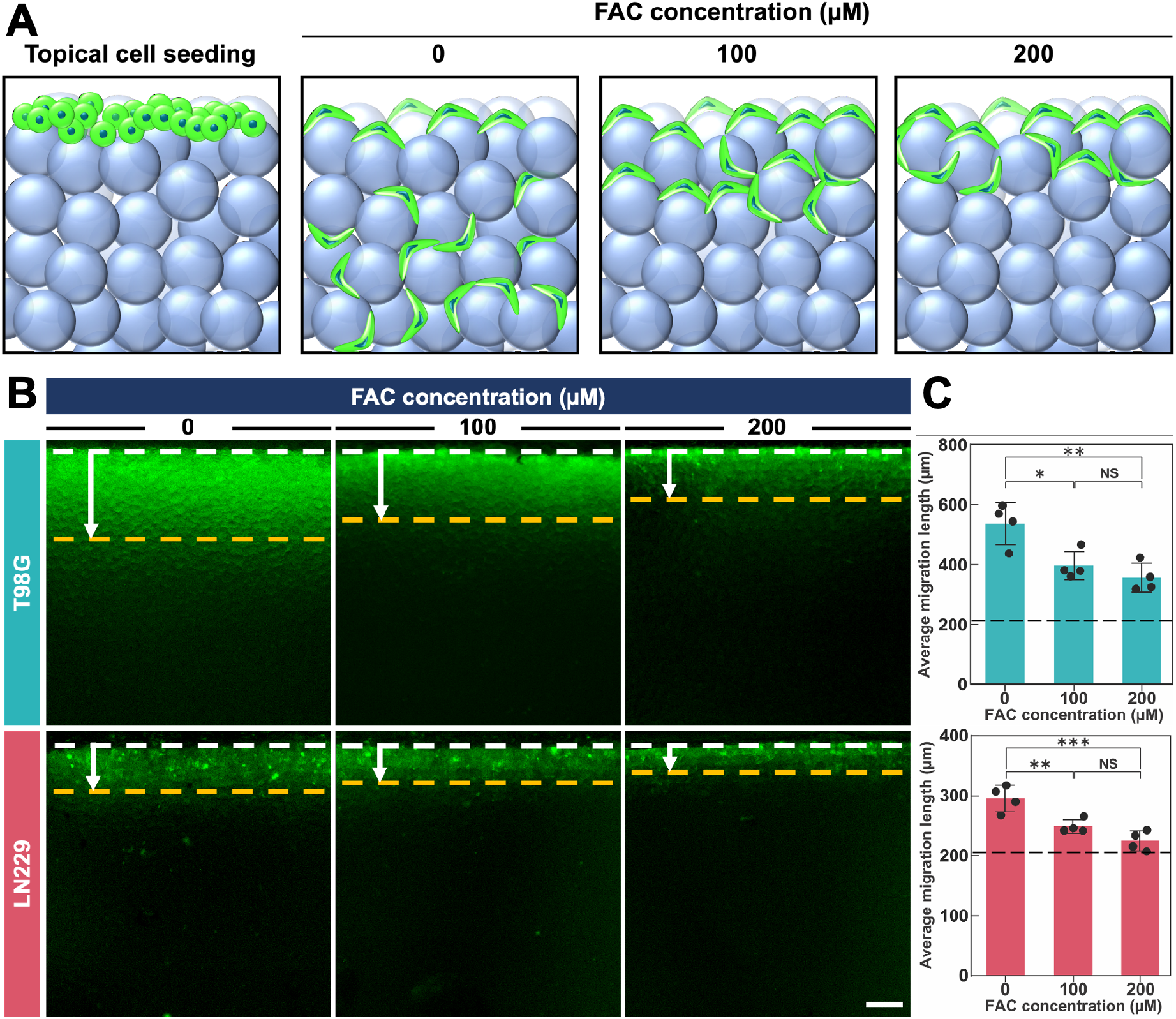
Effect of iron treatment on cancer cell migration in 3D granular hydrogel scaffolds. **(A)** Schematic illustration of cancer cell (T98G or LN229) topical seeding on 3D granular scaffolds, followed by cell culture in varying ferric ammonium citrate (FAC) concentrations (0, 100, or 200 μM). **(B)** Cross-sectional fluorescence microscopy images of T98G and LN229 cell lines migrated in the scaffold after 72 h of incubation. Adding FAC to the culture media resulted in significantly lower cell migration length. White and yellow dashed lines indicate the scaffold surface and average migration length, respectively (scale bar is 200 μm). **(C)** Average migration length at different FAC concentrations for T98G or LN229 cell lines (*n* = 4). The dashed lines show the average initial cell penetration length post topical seeding (~ 4 h) for all study groups pertinent to the T98G (212 ± 21 μm) or LN229 (207 ± 25 μm) cell lines (*n* = 3). The statistical analysis between average initial penetration length and average migration length for T98G cell line showed significant differences for all the study groups: 0 μM FAC (****p <0.0001), 100 μM FAC (**p < 0.01), or 200 μM FAC (*p <0.05). Similar analysis for the LN299 cell line showed significant differences for 0 μM FAC (****p <0.0001) and 100 μM FAC (*p < 0.05), and a nonsignificant difference for 200 μM FAC.

### 3.3 Iron at Tested Concentrations Does Not Reduce Cell Proliferation in T98G and LN229 Glioblastoma Cells

While we minimized any confounding due to cell proliferation in our wound healing assays by limiting the analysis to 24 hours and using relative wound density (RWD) as the measure, we decided to further verify that reduced cell proliferation due to iron-induced toxicity was not a confounder by directly assessing cell proliferation in T98G and LN229 cells at the iron concentrations used for the wound healing and 3D migration assays. In both T98G and LN229 cells, proliferation was not decreased even at 300 μM FAC as assessed by both optical confluence measurements (**Figure 3A, 3B**) or CyQuant™ cell proliferation assays (**Figure 3C, 3D**). In fact, in T98G cells, iron at concentrations of 100 and 200 μM appeared to promote proliferation as assessed by CyQuant™ cell proliferation assays (**Figure 3C**). We further verified that iron was not inducing cytotoxicity by measuring extracellular lactate dehydrogenase (LDH) levels in cells that were treated with iron. Treatment with up to 400 μM iron for 24 hours produced no significant increase in extracellular LDH in either cell line (**Figure 3E, 3F**), indicating that the iron concentrations used were not cytotoxic during the time periods where migration was assessed. These findings suggest that the inhibition of cell migration observed in the wound healing and 3D cell migration assays was specifically due to decreased cell migration and not simply decreased cell proliferation or increased cytotoxicity. Furthermore, our findings in the wound healing and 3D cell migration assays demonstrated a saturatable decrease in cell migration (i.e., both the addition of 200 μM and 300 μM appeared to produce a similar decrease in cell migration) whereas a dosedependent decrease in cell migration would be expected if iron-induced toxicity were playing a role.

**Figure 3.**
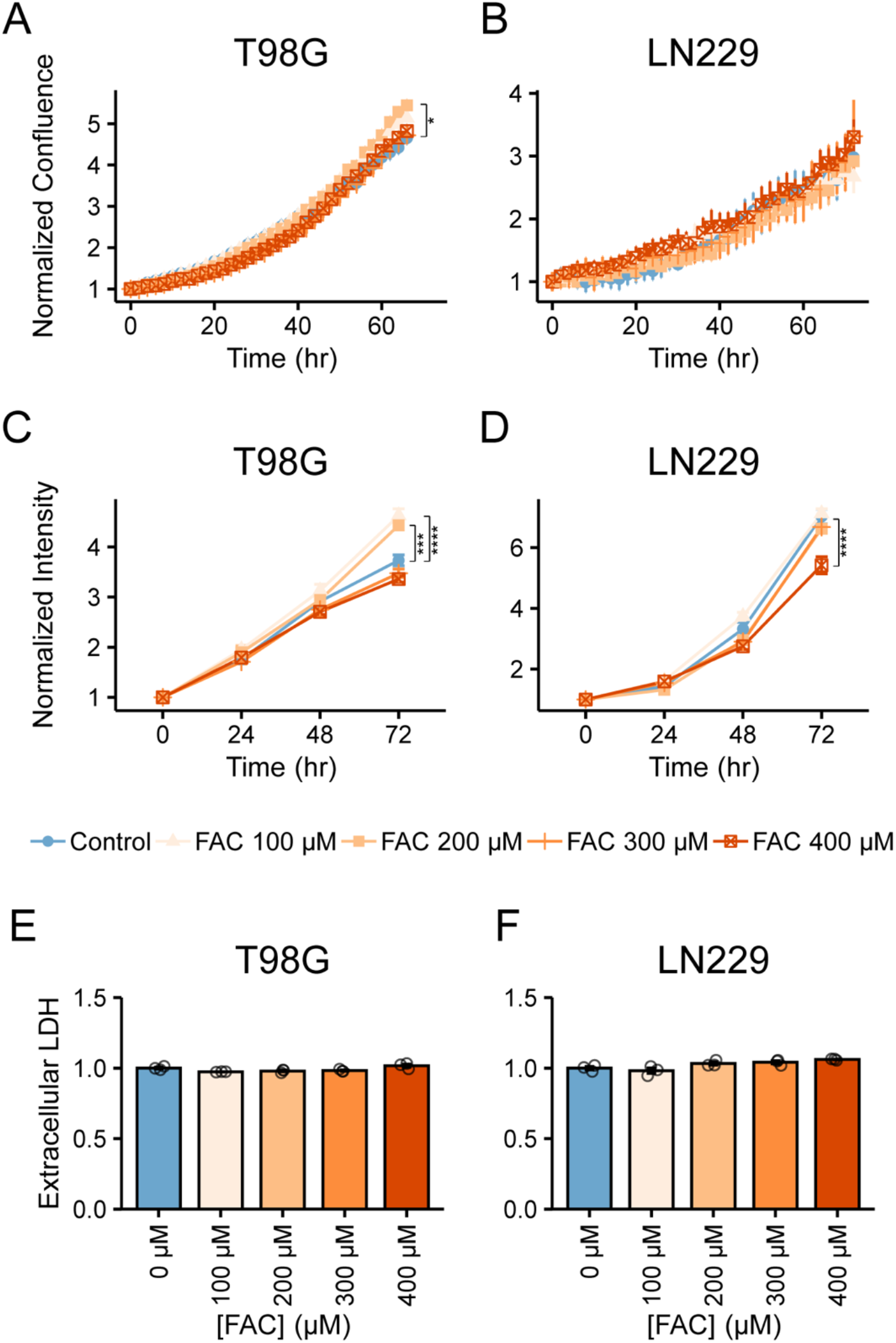
Iron does not inhibit cell proliferation or confluence in T98G or LN229 cells. **(A)** T98G or **(B)** LN229 glioblastoma cells were seeded into 96-well plates and allowed to proliferate over a period of three days in the presence of 0 – 400 μM ferric ammonium citrate (FAC). Optical confluence of the cells was measured as a function of time and appeared to be unaffected at the tested ferric ammonium citrate concentrations. **(C)** T98G or **(D)** LN229 cells were allowed to proliferate for 72 hours. Cell proliferation was measured at 24, 48, and 72 hours using a CyQuant® cell proliferation assay and found not to be reduced. Extracellular lactate dehydrogenase levels were not increased in **(E)** T98G or **(F)** LN229 cells after treatment with 0 – 400 μM ferric ammonium citrate for 24 hours. Results representative of at least n = 3 replicates. Student’s two-tailed t-test: *: p < 0.05, **: p < 0.01, ***: p < 0.001, ****: p < 0.0001.

### 3.4 Iron Reduces Cellular Polarization in T98G and LN229 Glioblastoma Cells

Since proper polarization of cells in the direction of migration is essential for efficient cellular locomotion, we employed a previously described cellular polarization assay to evaluate whether iron could be reducing cellular polarization [20,21]. Briefly, confluent monolayers of T98G or LN229 glioblastoma cells were scratched to generate a cell-free zone and cells were allowed to migrate into this zone for 6 hours. The cells were then fixed and stained with DAPI and anti-pericentrin to visualize the nucleus and the centrosomes, respectively. A cell was classified as correctly polarized if the centrosome fell within the quadrant of the cell facing the cell-free zone relative to the nucleus (**Figure 4A**). The number of correctly polarized cells in response to iron treatments was then quantified by an individual blinded to the experimental conditions. As hypothesized, the addition of iron in the form of 100 – 200 μM FAC or 50 – 100 μM hemin resulted in a significant decrease in the percent of correctly polarized cells (**Figure 4B – E**). These results show that in addition to reducing migratory capacity, iron inhibits the ability of glioblastoma cells to correctly polarize in the direction of migration.

**Figure 4.**
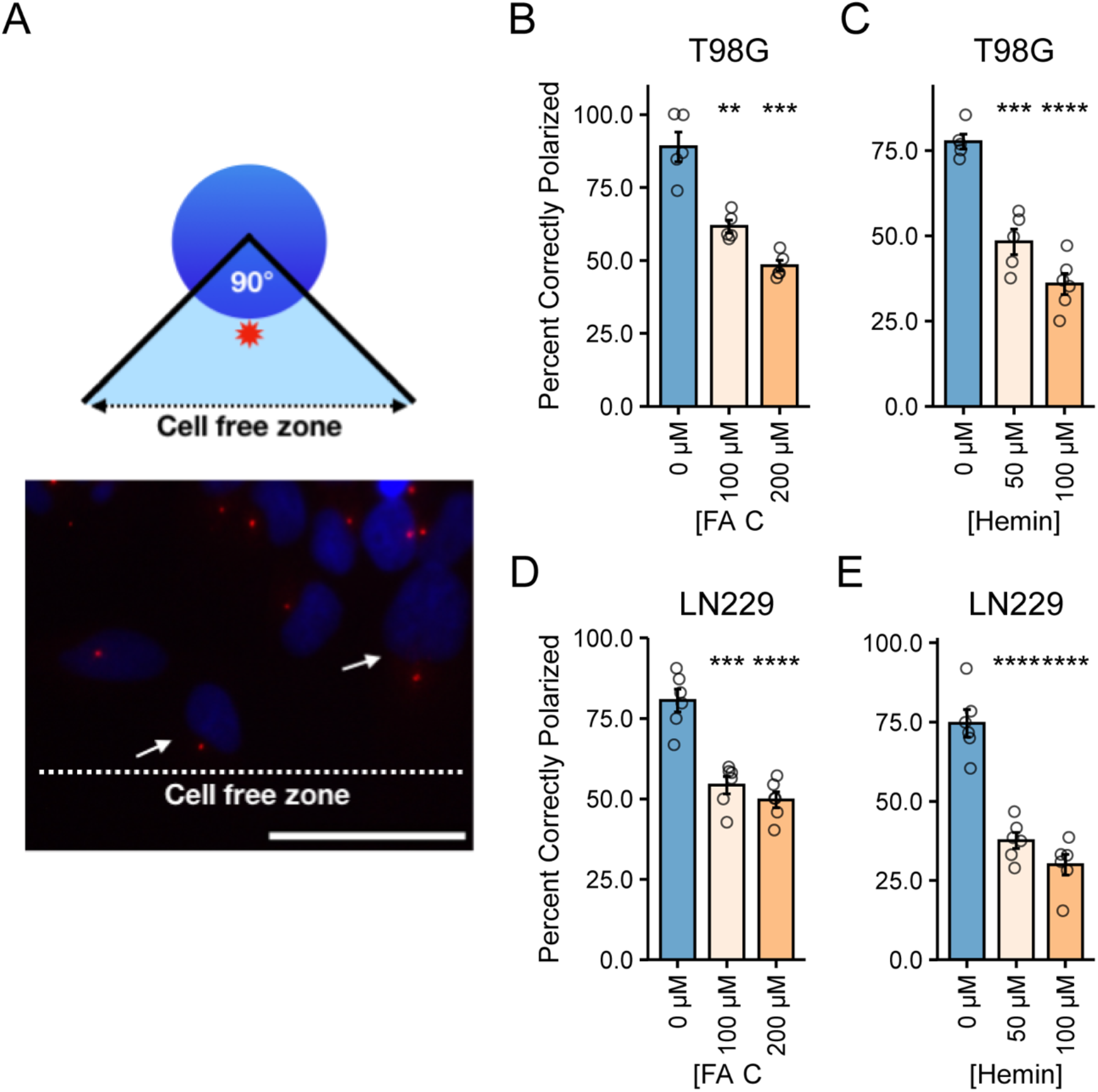
Iron reduces cell polarization in T98G and LN229 cells. A confluent monolayer of **(B, C)** T98G or **(D, E)** LN229 glioblastoma cells was scratched to generate a cell-free zone. Cells were then treated with 0 – 200 μM ferric ammonium citrate (FAC) or 0 −100 μM ferric chloride heme (Hemin) and allowed to migrate for 6 hours. Cells were fixed and stained with DAPI and anti-pericentrin to visualize cell polarity. Cells with pericentrin in the quadrant facing the cell-free zone relative to the nucleus were scored as correctly polarized (A), arrows point to correctly polarized cells. Results are representative of 50-90 cells for each condition scored over n = 4-6 replicates. Student’s two-tailed t-test: *: p < 0.05, **: p < 0.01, ***: p < 0.001, ****: p < 0.0001.

### 3.5 Iron Reduces Expression of Rho GTPase CDC42 in T98G and LN229 Glioblastoma Cells

CDC42 is a Rho GTPase that plays an essential role in cell migration by properly orienting cells and polarizing them in the direction of movement [11,12]. Since both cellular migration and polarization were reduced in response to iron, we sought to examine whether iron may alter expression levels of this Rho GTPase. Indeed, addition of iron in the form of 100 – 200 μM FAC or 50 – 100 μM hemin resulted in decreased expression of CDC42 in both T98G and LN229 glioblastoma cells (**Figure 5A-D**). One of the well described mechanisms of iron-mediated regulation of gene expression is through the iron-regulatory protein (IRP) - iron-responsive-element (IRE) system [26]. We first verified that our iron treatments resulted in increased cellular iron using SiRhoNox-1, a dye capable of labeling cellular ferrous iron (**Figure S1**). We then sought to determine whether the mRNA of CDC42 contained an IRE using the Searching for Iron Responsive Elements (SIREs) bioinformatic tool [27]. Analyzing CDC42 mRNA transcripts using SIREs revealed that a CDC42 mRNA splice variant (NM_001791.4) possessed an IRE-like sequence on the 3’ UTR which would be consistent with increased degradation of the mRNA in high iron conditions.

**Figure 5.**
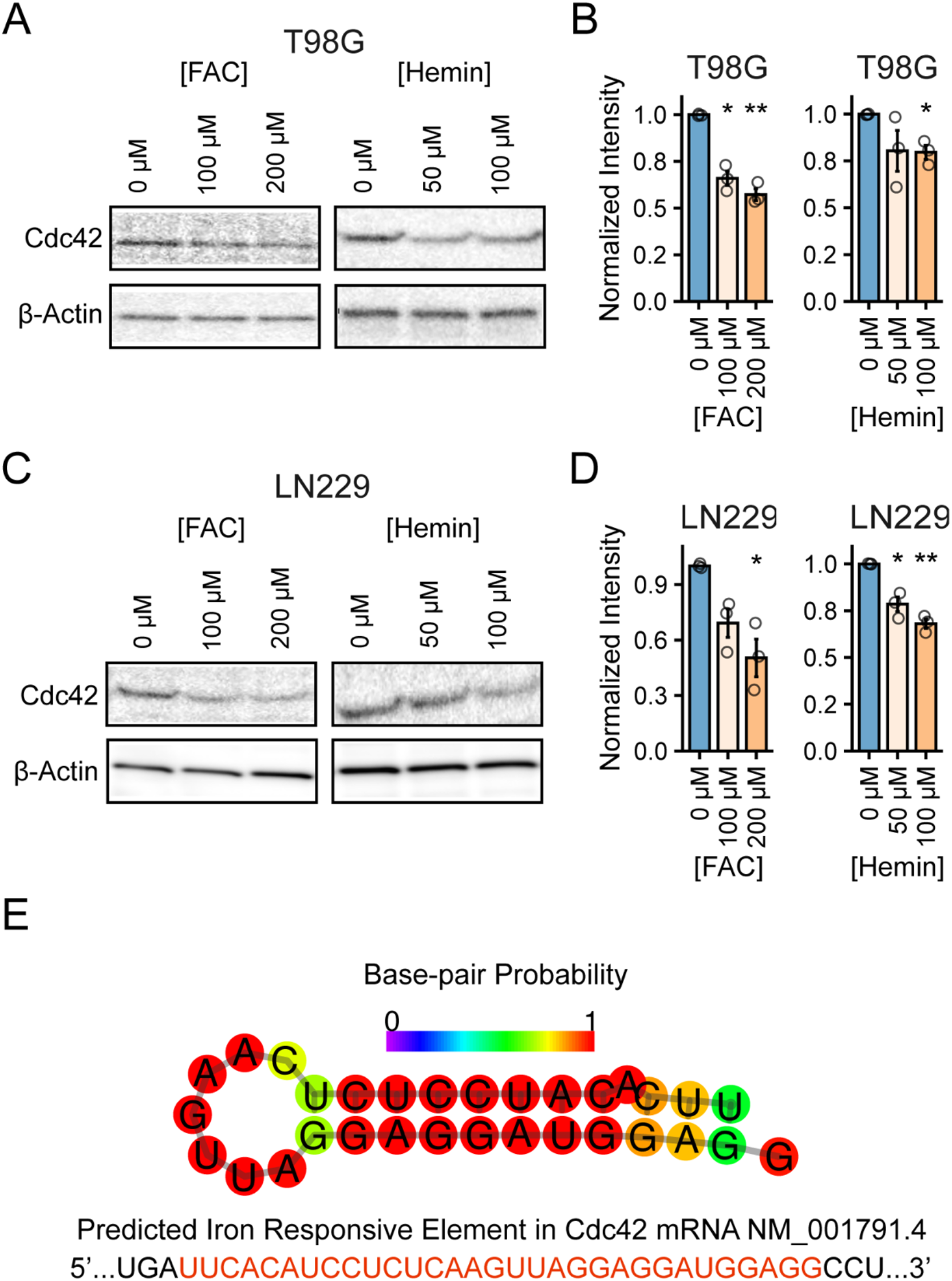
Iron decreases expression of CDC42 in T98G and LN229 cells. **(A)** T98G or **(C)** LN229 glioblastoma cells were treated with 0 – 200 μM ferric ammonium citrate (FAC) or 0 – 100 μM ferric chloride heme (hemin) for 6 hours and protein expression of CDC42 was measured by immunoblotting. Quantification from three independent experiments **(B, D)** showed that CDC42 expression was decreased in the iron treated cells. Results representative of n = 3 experiments. **(E)** Searching for Iron Responsive Elements (SIREs) was used to identify a potential iron-responsive element on the 3’ UTR of a CDC42 mRNA transcript. RNAFold through The Vienna RNA Websuite predicted a characteristic hairpin secondary structure on the IRE-like sequence identified by SIREs. Student’s two-tailed t-test: *: p < 0.05, **: p < 0.01.

SIREs utilizes multiple criteria including presence of a predicted loop structure, lack of mismatches, appropriate nucleotide pairing, and a predicted C8 bulge in the loop structure when determining whether a given mRNA sequence is likely to contain an IRE [27]. We further verified the loop structure of the predicted IRE sequence using RNAfold through the Vienna RNA Websuite [28–30] which also predicted the presence of a characteristic hairpin loop found on IREs (**Figure 5E**).

While the sequence for CDC42 mRNA NM_001791.4 is predicted to have an IRE, mismatches in the IRE were also predicted and a C8 bulge was not predicted, thus it remains to be determined whether the hypothesized IRE is functional *in-vitro and in-vivo*. Interestingly, the mRNA of CDC42 binding protein kinase alpha (CDC42BPA), a downstream effector of CDC42, has been experimentally confirmed to have an iron-responsive element [31] suggesting that this pathway may indeed be regulated through the IRP-IRE system.

### 3.6 Glioblastoma Cells at the Tumor Periphery Express Gene Signature Associated with Decreased Iron Stores

We investigated whether cells at the tumor periphery (i.e the cells spreading into healthy brain) differed from cells at the tumor core in terms of expression of iron-related genes in human glioblastoma patient samples. We downloaded single-cell RNA-seq data from a previously published dataset (located at gbmseq.org) [22] and compared expression of key iron regulatory genes between cells at the tumor periphery to those at the core (**Figure 6A**). Cells at the periphery expressed higher levels of *TFRC*, the gene for transferrin receptor (**Figure 6B**). Transferrin receptor-mediated uptake of iron represents a primary means of iron uptake in mammalian cells. Expression of *TFRC* is strongly regulated by cellular iron stores with cells with lower levels of iron upregulating *TFRC* to increase iron uptake and achieve homeostasis [32]. Cells at the periphery also expressed lower levels of *FTH1* and *FTL* (**Figure 6C, 6D**), the genes for H- and L-ferritin, respectively. Ferritin is an essential iron storage protein composed of an H- and L-subunits. Expression of *FTH1* and *FTL* have been demonstrated to be positively associated with cellular iron content with cells with higher iron stores having higher levels of ferritin to safely manage the iron and not induce excessive oxidative stress [33]. Higher gene expression of *FTH1* and *FTL* and lower expression of *TFRC* in cells at the tumor core is consistent with cells at the core (i.e cells that are not spreading into healthy brain) having higher levels of iron and those at the periphery (i.e cells that are spreading into healthy brain) having lower iron content.

**Figure 6.**
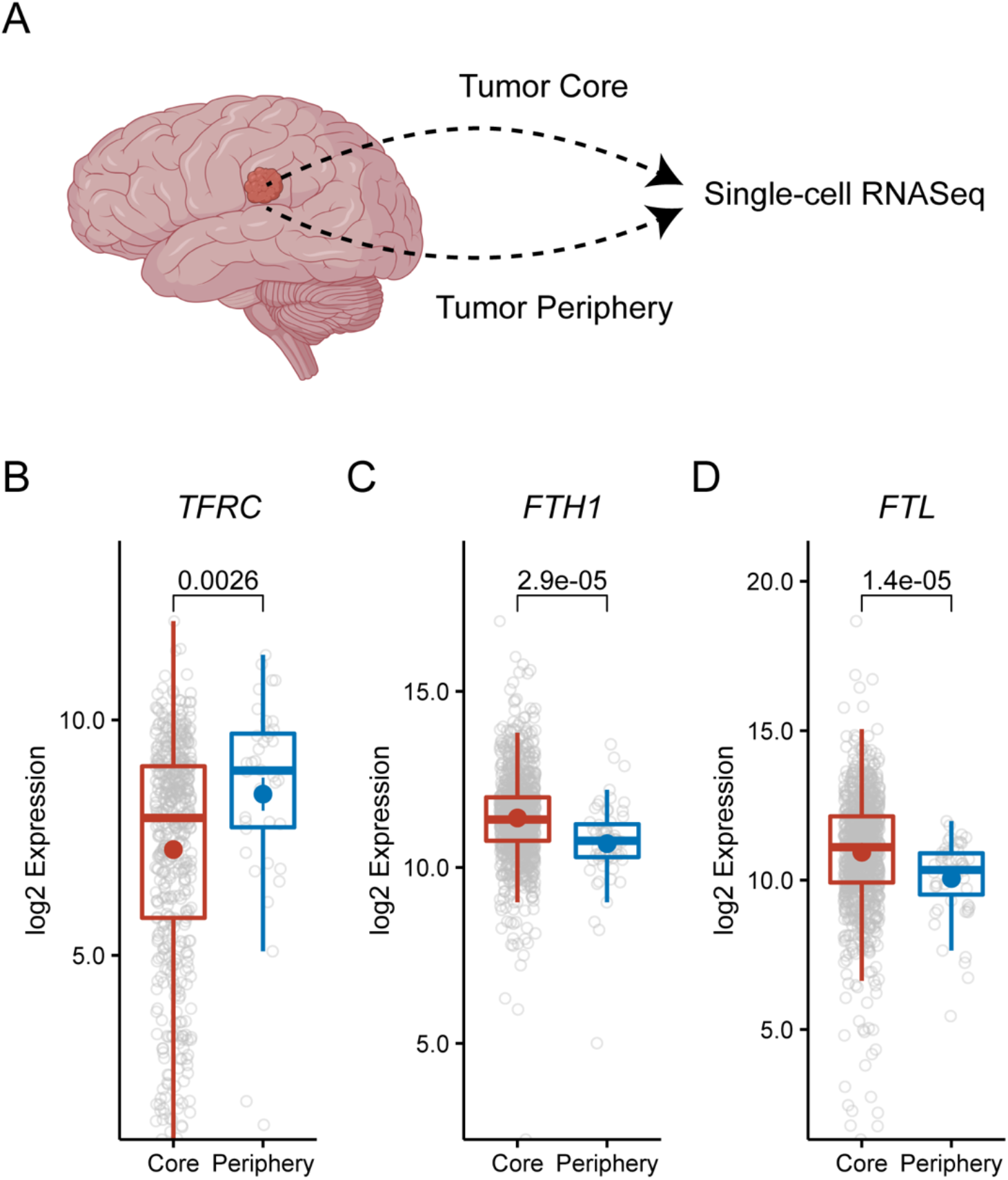
Glioblastoma cells at the tumor periphery have gene signature associated with lower iron levels. **(A)** A single cell RNA-Seq dataset (GSE84465) characterizing 1091 neoplastic cells split between the tumor core and periphery from four glioblastoma patients was analyzed for expression of markers associated with cellular iron status, including **(B)** transferrin receptor (*TFRC*), **(C)** H-ferritin (*FTH1*), and **(D)** L-ferritin (*FTL*). Cells at the tumor periphery had increased expression of *TFRC* and decreased expression of *FTH1* and *FTL*, a gene signature consistent with cells that have lower iron stores. P-values are derived from two-tailed Student’s t-test. Created with BioRender.com.

## 4. Discussion

Historically, iron has been thought to promote tumor growth because it directly supports the high rate of proliferation of cancer cells. While iron’s ability to promote cell proliferation at sub-toxic concentrations has indeed been well documented, other impacts of iron on cancer cell biology, including cell migration, have been less thoroughly characterized. In recent years the double-edged nature of iron in cancer biology has begun to be appreciated with discoveries such as ferroptosis, a form of iron-dependent cell death [34,35], demonstrating that iron may potentially have favorable roles in cancer treatment. Here we add to the literature describing iron’s numerous impacts on cancer cell biology by demonstrating that iron alters the ability of glioblastoma cells to migrate – a deadly feature that enables these cells to diffusely spread into healthy brain and generate recurrent disease.

The results presented here show that iron inhibits cell migration in T98G and LN229 human glioblastoma cells. The addition of deferoxamine, an iron chelator, was able to largely restore migratory capacity, confirming that iron was responsible for our findings. Interestingly, the iron-induced inhibition of migration displayed a saturatable response with concentrations above 200 μM iron displaying similar degrees of inhibition. This saturatable response indicates that glioblastoma cells have an intrinsic regulator that signals the cells to limit migration when iron is available.

Since treatment-induced effects on proliferation can potentially confound studies examining cell migration, a thorough investigation into iron’s effect on proliferation was carried out, revealing that cell proliferation was not reduced at the concentrations used in the migration assays. If anything, proliferation was stimulated in T98G cells at 100 μM iron, further confirming that the observed effects are indeed due to reduced migration. Additionally, toxicity brought about by iron would be expected to exhibit a dose-dependent response whereas we observed a saturatable response with iron concentrations above 200 μM producing similar degrees of migration inhibition.

Mechanistically, the addition of iron reduced expression of CDC42, a Rho GTPase essential to cell migration, in our glioblastoma cells. CDC42 establishes cellular polarity during migration by remodeling the cellular cytoskeleton and orienting centrosomes toward the direction of movement [12]. Numerous studies have implicated CDC42 in the migration of various types of cancer cells [36–39]. Interestingly, SIREs, a bioinformatic tool developed to detect iron responsive elements (IREs) on mRNA transcripts [27], predicted the presence of an IRE on the 3’ UTR of a CDC42 mRNA transcript. IREs are short hairpin-loop regulatory sequences found in untranslated regions (UTRs) of mRNAs involved in iron-related processes such as cell metabolism, iron storage, and hypoxia signaling [26]. In low iron conditions, iron-regulatory protein 1 (IRP1) and iron-regulatory protein 2 (IRP2) can bind to IREs on target mRNAs and influence their stability while high iron conditions prevent binding of IRPs to IREs. Binding of IRP1/IRP2 to an IRE on the 3’ UTR of an mRNA leads to stabilization and increased translation of the mRNA while binding of IRP1/IRP2 to an IRE on the 5’ UTR of an mRNA leads to target degradation [26]. Thus, the SIREs predicted IRE on the 3’ UTR of CDC42 mRNA would be consistent with our findings of iron-induced reduction in CDC42 expression. However, we do note that SIREs also predicted mismatches and did not predict a C8 bulge typically found on the secondary structure of IREs in the CDC42 transcript, thus it remains to be determined whether the IRE on CDC42 mRNA is functional. Nonetheless, we verified that the iron-induced decrease in CDC42 expression in these cells was also accompanied by a functional decrease in cellular polarization. To the best of our knowledge, this is the first demonstration of iron’s ability to impact cellular polarization in human glioblastoma cells.

Our cell culture data would predict that neoplastic cells in the tumor periphery would have less iron as they would be expected to be enriched in the population of cells migrating away from the core. We thus analyzed a single-cell RNASeq dataset of human glioblastoma samples [22] that revealed cells at the tumor periphery express higher levels of transferrin receptor (*TFRC*) and lower levels of H-ferritin (*FTH1*) and L-ferritin (*FTL*). Expression of *TFRC* is negatively regulated by cellular iron levels with lower iron levels inducing expression of *TFRC*. Similarly, *FTH1* and *FTL* are also regulated by cellular iron levels with higher iron levels inducing expression of these iron-storage proteins [33,40]. Thus, higher levels of expression of *TFRC* and lower levels of expression of *FTH1* and *FTL* for neoplastic cells in the tumor periphery are consistent with our cell culture findings that migrating cells have lower levels of cellular iron and that the cells with higher iron may be less migratory and thus less likely to leave the tumor core.

An explanation for the ability of iron to inhibit glioblastoma cell migration lies in the possibility that it may be advantageous for a cell to reduce motility in a favorable environment (i.e., remain in an environment where there is sufficient iron and nutrients to proliferate). Along these lines, the “go or grow” hypothesis postulates that cells may switch between migratory and proliferative states such that proliferating cells have decreased motility, although recent studies have cast doubt on the validity of the “go or grow” hypothesis, at least in melanoma cells [41].

Nonetheless, our findings raise the question of whether iron could play a role in glioblastoma treatment. While it has been documented to promote cancer cell proliferation at sub-toxic concentrations, iron’s impact on clinical endpoints such as overall survival among glioblastoma patients is less clear. Several clinical studies have demonstrated that anemic glioblastoma patients (i.e., patients with absolute or functional iron deficiency anemia) had poorer clinical outcomes than those that were not anemic [42–45]. Similarly, other studies have found that the administration of iron in the form of iron oxide nanoparticles improved survival outcomes in mouse models of glioma [46] and breast cancer [47]. Our findings may potentially help explain an underlying process contributing to these seemingly counterintuitive outcomes.

Clinical endpoints and response to therapeutic intervention depend on more than just cancer cell proliferation rate. For example, a tumor that is proliferating faster but not disseminating into healthy brain will be more amenable to complete surgical resection and more responsive to DNA alkylating agents, potentially conferring a better patient outcome than a tumor that proliferates slowly and disseminates widely into the brain. Strategies aimed at limiting the spread of glioblastoma cells into healthy brain, potentially through manipulating iron metabolism, could yield improvements to patient outcomes. Furthermore, iron may impact other features of glioblastoma tumor biology such as induction of hypoxic responses [48–50], immune function [51], and susceptibility to ferroptosis [34,35] that may offset its contribution to cell proliferation. Our findings shed light on the largely uncharacterized role of iron in glioblastoma cell migration and demonstrate the need for further investigation into the role of iron in cancer cell and tumor microenvironment biology.

## Acknowledgements

The authors would like to thank the members of the Connor, Lathia, Wang, and Sheikhi labs for helpful discussions and technical assistance. J.R.C and G.S would like to acknowledge funding received from the Department of Neurosurgery at the Penn State College of Medicine, the Tara Leah Witmer Memorial Fund, and the National Institutes of Health (F30CA250193 to G.S and P01CA245705 to J.R.C). The contents of this paper are solely the responsibility of the authors and do not necessarily represent the official views of the National Institutes of Health. Amir Sheikhi would like to acknowledge the funding received from the Meghan Rose Bradly Foundation.

## Supplementary Information

### 1. Supplementary Methods

#### 3D Hydrogel Scaffold Synthesis

##### Materials

Gelatin (Type A from porcine skin), methacrylic anhydride, and photoinitiator (lithium phenyl-2,4,6-trimethylbenzoylphosphinate, LAP) were purchased from Sigma (MO, USA). Dulbecco’s phosphate-buffered saline (DPBS) was obtained from Gibco (MA, USA), CellTracker™ Green CMFDA Dye was purchased from Invitrogen (MA, USA). Dialysis tubing (6-8 kDa molecular weight cutoff membranes) was obtained from Spectrum Laboratories (NJ, USA). Vacuum filtration unit (pore size = 0.20 μm) was purchased from VWR (PA, USA). Pico-surf™ and Novec 7500™ oil were obtained from Sphere Fluidics (Cambridge, UK) and 3M (MN, USA), respectively. 1H,1H,2H,2H-Perfluoro-1-octanol was purchased from Alfa Aesar (MA, USA). Twenty four-well plate non-treated cell culture plate was obtained from Cellstar (Greiner Bio-One, Austria).

##### GelMA synthesis

GelMA was synthesized as previously described (15, 23). In summary, 20 g of gelatin (Type A from porcine skin) was fully dissolved in 200 mL of DPBS at 50 °C, to which 2.5 mL of methacrylic anhydride was added dropwise and allowed to react for 2 h under stirring. The reaction was stopped by adding 400 mL of 50 °C DPBS. The solution was dialyzed with 6-8 kDa molecular weight cutoff membranes against a stirring 40 °C ultra-pure Milli-Q® water (Millipore Corporation, MA, USA) for 10 days to remove methacrylic anhydride and any other byproduct. The solution obtained from dialysis was sterile vacuum filtered using a 0.20 μm vacuum filtration system and was frozen at −80 °C, followed by lyophilization for 3 days.

##### GelMA microgel fabrication

To fabricate GelMA microgels, a step emulsification microfluidic device was fabricated and used according to a previously established protocol (15, 17). For the dispersed phase, freeze-dried GelMA was dissolved in 40 °C DPBS containing 0.1% w/v photoinitiator (LAP) at a concentration of 20 mg mL^-1^. The continuous phase consisted of Pico-surf™ (2% v/v) in Novec 7500™ oil. Two syringe pumps (PHD 2000, Harvard Apparatus, MA, USA) were used to inject the dispersed and continuous phases separately into the device at a flow rate of 100 and 200 μL min^-1^, respectively, while a space heater maintained the system temperature at 35-40°C. To create physically crosslinked GelMA microgels, the droplet suspension in oil was stored at 4 °C, while protected from any light.

##### Scaffold preparation

To remove the oil and surfactant from the GelMA microgel suspension, Novec 7500™ oil supplemented with 1H,1H,2H,2H-Perfluoro-1-octanol (20% v/v) was added to the droplet suspension (1:1 volume ratio) and vortexed for 5 s. The suspension was then centrifuged at 325 ×**g** for 15 s. DPBS containing 0.1% w/v of LAP at 4 °C was added to the washed suspension and vortexed for 5 s, followed by two times centrifugation at 325 ×**g** for 15 s. During the washing process, a cold-water bath was used to maintain the suspension at 4 °C. After discarding the supernatant, a positive displacement pipette (Microman E M100E, Gilson, OH, USA) was used for pipetting the microgel suspension on a laser-cut acrylic mold (diameter = 10 mm and thickness = 3 mm). Then the molded microgel suspension was exposed to light (wavelength of 395-405 nm and intensity of 15 mW/cm^2^ for 1 min), to undergo covalent bond-mediated microgel assembly.

### 2. Supplmentary Figures

**Figure S1.**
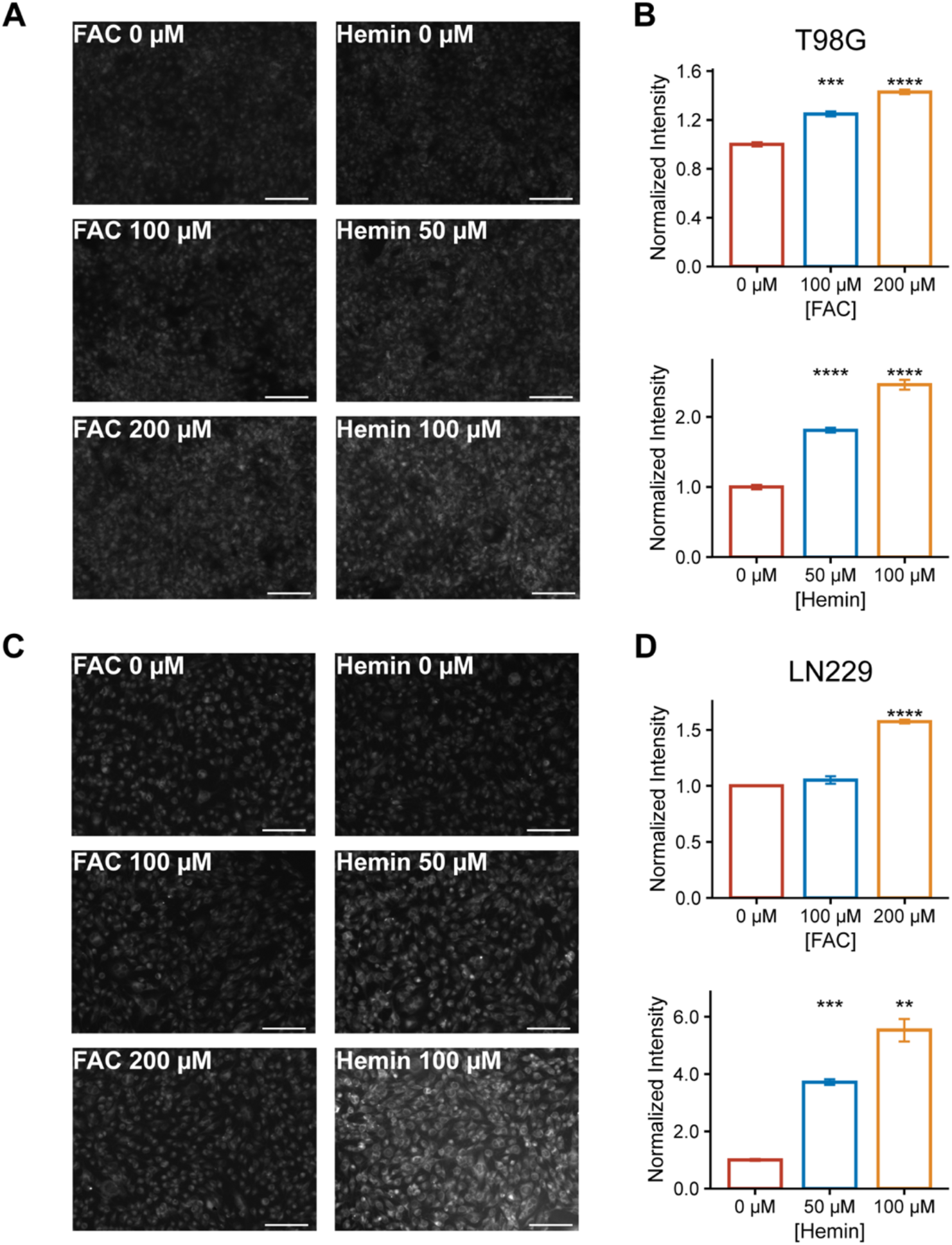
Addition of ferric ammonium citrate (FAC) or ferric chloride heme (Hemin) increases labile iron pool in T98G and LN229 cells. (A, B) T98G or (C, D) LN229 cells were treated with 0 – 200 μM ferric ammonium citrate or 0 – 100 μM ferric chloride heme and the labile iron pool was measured using siRhoNox-1, a dye that specifically detects Fe^2+^. Results are representative of images quantified from n = 3 replicates. Student’s two-tailed t-test: *: p < 0.05, **: p < 0.01, ***: p < 0.001, ****: p < 0.0001.

